# Long-Term Grazing Drives Compositional Shifts in Root-Associated Microbial Communities of Desert Steppe Plants

**DOI:** 10.64898/2026.08.24.746874

**Authors:** Aimin Zhu, Fan Jiang, Shuai Luo, Zhichun Yan, Xu Cheng, Guodong Han, Ton Bisseling

## Abstract

Grassland microbial communities are central to mediating ecosystem function and stability, yet how long-term grazing reshapes root-associated microbiomes across contiguous soil-root habitats remains poorly understood. This limits our ability to identify robust microbial bioindicators for grassland health monitoring. In this study, we investigated the community assembly and functional variation of root-associated microbiomes of *Stipa breviflora*, a dominant perennial clonal grass in desert steppes, across a 17-year continuous grazing experiment with four grazing intensity treatments (no grazing, light, moderate, and heavy grazing). We show that grazing intensity induces niche-specific restructuring of microbial communities, with the most profound compositional and functional shifts occurring in the rhizosphere, followed by root endophytic compartments and bulk soil. Light and moderate grazing significantly enriches the phylum Bacillota in rhizosphere and endophytic compartments, whereas the genus *Pseudomonas* dominates ungrazed grassland soils and is markedly depleted under grazing conditions. Microbial community responses to grazing follow a unimodal intermediate disturbance pattern, with moderate grazing triggering the strongest microbial community differentiation, enhanced microbial network connectivity and modularity, and the highest abundance of grazing-responsive microbial biomarkers. Notably, grazing-induced microbial community variation is decoupled from intraspecific phenotypic changes in *S. breviflora*. Our findings demonstrate that long-term grazing acts as a strong selective filter partitioning core beneficial microbial taxon, establishing Bacillota and *Pseudomonas* as complementary bioindicators for evaluating desert steppe ecosystem health. This study advances the understanding of plant-microbe interactions under anthropogenic disturbance and provides microbiome-based insights for sustainable grassland management.

## Introduction

Grasslands represent one of the largest terrestrial biomes, covering approximately 26% of Earth’s surface and providing critical ecosystems services including climate regulation, carbon sequestration, and biodiversity conservation (Van den Berg et al., 2011; Ma et al., 2018; Zhao et al., 2020). As the most prevalent anthropogenic disturbance in grassland ecosystems, livestock grazing profoundly modulates plant community structure, soil physicochemical properties, and belowground microbial assemblages via herbivory, soil trampling, and animal nutrient return (Liu et al., 2018; Che et al., 2019, Han et al., 2020). Livestock grazing effects are strongly context-dependent, varying with grazing intensity, livestock type, and local soil and vegetation characteristics (Zhang et al., 2018; Bakhshi et al., 2020).Belowground microbial communities are pivotal drivers of grassland ecosystem function, regulating soil nutrient cycling, organic matter decomposition, and plant growth promotion via intimate plant-microbe interactions (Kreuzer et al., 2004; Bardgett et al., 1996).

Soil microorganisms are pivotal to grassland functioning, regulating nutrient cycling, organic matter turnover, and plant growth through processes such as decomposition of organic material and interactions with the roots (Kreuzer et al., 2004). Mounting evidence indicates that grazing intensity differentially shapes soil microbial diversity and activity. Moderate grazing generally enhances root exudation and soil nutrient input, stimulating microbial diversity and metabolic activity, while heavy grazing induces soil compaction, organic matter loss, and microbial community degradation (Steenwerth et al., 2002; Qiao, 2008). Rhizosphere and endophytic compartment are host the root-associated microbiomes, with taxa affiliated with Pseudomonadaceae and Rhizobiaceae widely recognized for enhancing plant growth and stress tolerance (Berlemont et al., 2013; Yan et al., 2017; Wang et al., 2019). Although plant endophytic bacteria contribute substantially to host stress resistance and growth regulation, their niche-specific responses to long-term grazing remain unstudied.

Desert steppes are fragile, water-limited grassland ecosystems highly vulnerable to anthropogenic disturbance, yet the mechanisms linking long-term grazing, niche-specific microbial community assembly, and ecosystem stability remain poorly characterized. Existing grazing microbiome studies predominantly focus on bulk soil or rhizosphere communities, with limited systematic comparisons across bulk soil, rhizosphere, and root endophytic compartments (Hawkes et al., 2015; Singh et al., 2015; Fierer, 2017). Furthermore, the functional responses of ecologically critical microbial taxa—including stress-resistance promoting Bacillota and plant-beneficial *Pseudomonas*—to chronic grazing disturbance remain elusive (Lupwayi et al., 2024). This critical knowledge gap hinders our mechanistic understanding of grazing-induced grassland degradation and limits the development of microbiome-based ecosystem monitoring strategies.

In this study, we leveraged a 17-year continuous grazing field experiment in a typical Inner Mongolian desert steppe, focusing on *Stipa breviflora*, the dominant constructive perennial grass species of the region. We systematically investigated grazing intensity effects on microbial community diversity, composition, co-occurrence networks, and biomarker taxa across three interconnected root-associated habitats (bulk soil, rhizosphere soil, and root endophytic compartments). We specifically tested two core hypotheses: (1) long-term grazing induces niche-dependent shifts in root-associated microbial community structure and diversity, with divergent responses across soil-root continuum habitats; (2) distinct microbial taxa (Bacillota and *Pseudomonas*) serve as differential bioindicators for low and high grazing disturbance, mediating ecosystem adaptive responses. By integrating niche-specific community profiling and taxonomic abundance analysis, our study elucidates the assembly rules of root-associated microbiomes under long-term grazing and provides robust microbial biomarkers for desert steppe ecosystem health assessment and sustainable management.

## Materials and Methods

### Study site and experiment design

This study was conducted at the long-term grazing experimental demonstration base of the Inner Mongolia Academy of Agriculture and Animal Husbandry Sciences, located in Siziwang Banner, Inner Mongolia, China (41 ° 47’17 “N, 111 ° 53“46” E, elevation 1,450 m). The grazing experiment was established in 2004 on a 50 hectares natural desert steppe, and field sampling was performed in May 2018 after 17 consecutive years of grazing treatment. The experiment adopted a randomized complete block design with three biological blocks and four grazing intensity treatments: no grazing (Control), light grazing (LG, 0.91 sheep ha), moderate grazing (MG, 1.82 sheep ha), and heavy grazing (HG, 2.71 sheep ha). Each experimental plot covered 4.4 ha, generating a total of 12 experimental units. Grazing was implemented annually from June to November using 2-year-old local sheep, with daily grazing duration from 6:00 to 18:00. Livestock were provided with sufficient drinking water and supplementary salt to maintain normal growth during the grazing period.

### Microbial sample collection

Microbial samples were collected along the central axis of each plot at 50 m intervals, with four sampling points per plot. At each sampling point, bulk soil (0–20 cm depth) and healthy *S. breviflora* root systems with attached rhizosphere soil were collected. Rhizosphere soil was obtained by gently shaking root systems to separate tightly adherent soil particles. Whole root systems and rhizosphere soil samples were placed in sterile plastic bags, transported to the laboratory in insulated cooling boxes, and stored at −20 °C for subsequent microbial DNA extraction. A total of 48 bulk soil samples, 144 rhizosphere soil samples and 144 root endosphere samples were initially collected. After DNA extraction and amplicon sequencing, 32 unqualified samples with low sequencing quality were excluded, yielding a final dataset of 304 valid samples for downstream bioinformatic and statistical analyses.

### DNA extraction and high-throughput PCR amplification

Microbial genomic DNA was extracted from bulk soil and rhizosphere soil samples using the MOBIO DNeasy PowerSoil Kit (12888-100). Root endosphere microbial DNA was extracted using the FastDNA Spin Kit for Soil following established protocols (Schneijderberg et al., 2020; Zhu et al., 2022). The V3 –V4 hypervariable region of the bacterial 16S rRNA gene was amplified via PCR using the universal primer pair 515F (5′ -GTGCCAGCMGCCGCGGTAA-3′) and 806R (5′-GGACTACHVGGGTWTCTAAT-3′). Purified PCR products were sequenced on the Illumina HiSeq platform (Beijing Novogene Bioinformatics Technology Co., Ltd.) for high-throughput amplicon sequencing.

### Amplicon sequencing and bioinformatic processing

Paired-end sequencing (2 × 250 bp) was performed on the Illumina HiSeq 2500 platform. Raw sequencing reads were quality-trimmed and filtered using fastp v0.24.0 with parameters -q 20 -l 100, retaining clean reads with an average error rate < 0.01 and read length ≥ 100 bp (Chen et al., 2018). High-quality clean reads were processed using the QIIME 2 v2023.2 pipeline (Bolyen et al., 2019). Amplicon Sequence Variants (ASVs) were resolved using the DADA2 algorithm with default parameters (Callahan et al., 2016).

Taxonomic annotation of ASVs was performed using a naive Bayesian classifier against the SILVA v138.1 reference database. Sequences annotated as chloroplasts and mitochondria were removed to eliminate eukaryotic contamination. ASVs with fewer than 25 total sequences or detected in fewer than five samples were filtered out to reduce sequencing noise, resulting in 4231 valid ASVs for subsequent analyses.

### Statistical analysis

All statistical analysis were performed in R software (v4.4.1). Linear mixed-effects models (LMMs) were implemented separately for each niche to analyze the effects of grazing intensity on alpha diversity and the relative abundances of individual microbial taxa. For both types of models, grazing intensity was included as a fixed effect, and the experimental block was incorporated as a random intercept effect to account for spatial dependencies. All models were fitted using the `lme4` package in R. For alpha diversity, model-estimated marginal means and their 95% confidence intervals for each grazing level within a niche were computed and compared using the `emmeans` package. For taxon-level analyses, effect sizes for grazing intensity were extracted, and p-values were adjusted for multiple comparisons across all tested taxa using the Benjamini–Hochberg false discovery rate (FDR) procedure, with an FDR-corrected p-value < 0.05 deemed statistically significant.

To calculate the alpha and beta diversity, the sequence of all samples was flattened to a uniform data volume. Because the sequence number in the root endosphere is relatively low compared to that from other compartments (Supplementary Table S2), we separately rarefied the root endosphere samples (11,141 sequences) and those from the other compartments (32,301 sequences), using the minimum sequence count within each group as the rarefaction depth.

For each sample, we computed alpha diversity metrics, including Shannon and Chao1 indices, using “vegan” package. We compared these metrics among different grazing intensity treatments using “multcompView” package.

To evaluate overall differences in bacterial community composition, we performed principal coordinate analysis (PCoA) based on Bray–Curtis dissimilarities. Separate ordinations were generated for each sampled niche (bulk soil, rhizosphere soil, and root endosphere) to visualize the effect of grazing intensity on community structure. The influence of grazing intensity, compartment niche, and their interaction on microbial community dissimilarity was tested using permutational multivariate analysis of variance (PERMANOVA) via the adonis function in the vegan package in R.

To identify potential microbial taxa that serve as biomarkers distinguishing grazing treatments, we used Linear Discriminant Analysis Effect size (LEfse). Taxa with an LDA effect size greater than 2.0 (*P* < 0.05) were considered significant. We additionally applied the Wilcoxon signed-rank test to to compare the relative abundance of taxa between control samples and those under each grazing intensity across multiple taxonomic levels. P-values from these pairwise comparisons were adjusted for multiple testing using the Benjamini–Hochberg false discovery rate (FDR) method, with an adjusted significance threshold of *P* < 0.05.

## Results

### Grazing intensity modulates phenotypic traits of *Stipa breviflora*

We quantified key growth traits of *S. breviflora* across all grazing intensity treatments to characterize plant phenotypic responses to long-term grazing (**Figure 1**). Relative to the ungrazed control, moderate grazing significantly increased plant height (mean = 20.0 cm), while light and heavy grazing induced a non-significant reduction in plant height. Vegetation coverage exhibited no obvious variation under light and moderate grazing, but increased notably under heavy grazing. Plant density and aboveground biomass displayed consistent increasing trends across all grazing treatments, with moderate grazing yielding the highest values for both indicators. Collectively, moderate grazing optimized the growth performance of *S. breviflora*, although all phenotypic differences relative to the control did not reach statistical significance.

**Figure 1.**
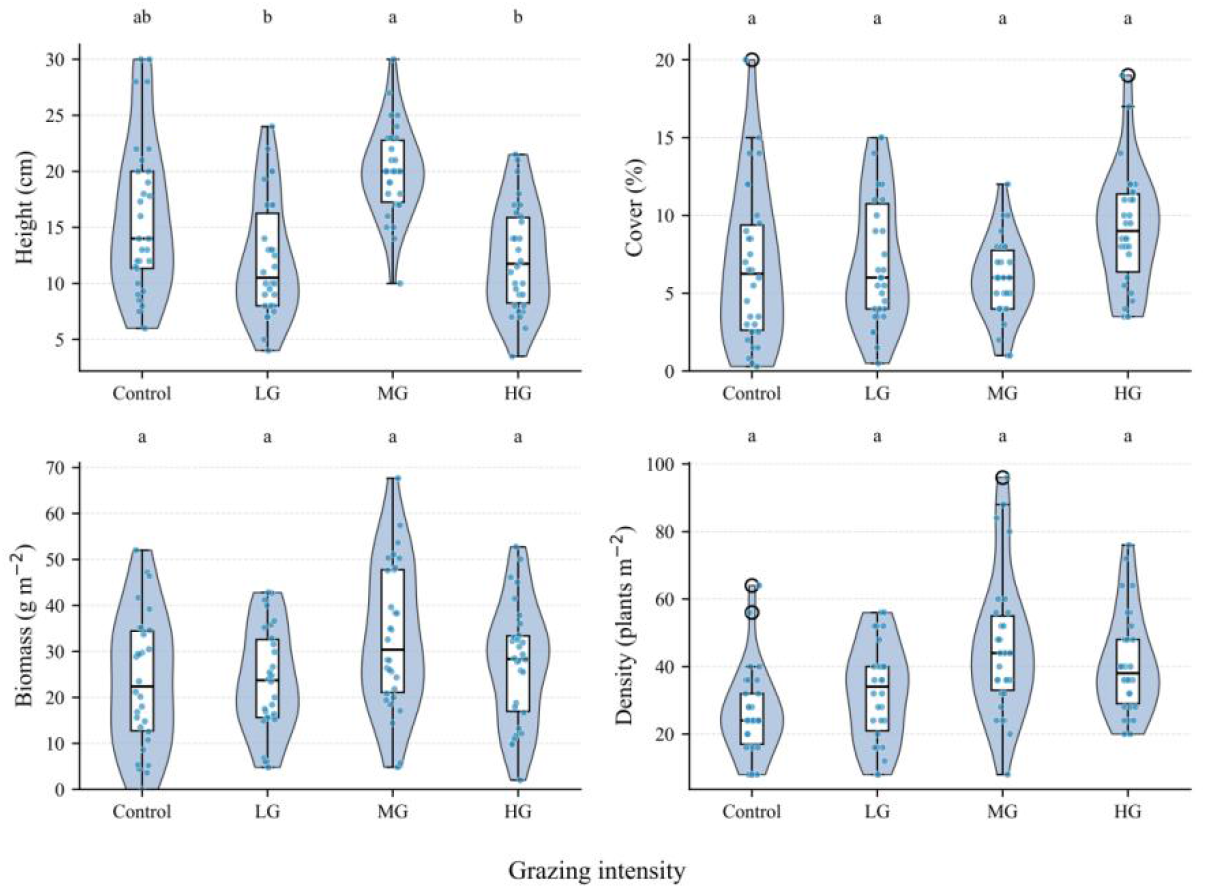
Effects of different grazing intensities on the height, cover, density, and biomass of *Stipa breviflora*. The different lowercase letters in the figure indicate significant differences at the *P*-value < 0.05. Control, no grazing; LG, light grazing treatment; MG, moderate grazing treatment; HG, heavy grazing treatment.

### Grazing intensity reshapes niche-specific microbial community diversity and structure

To further investigate the effects of different grazing intensities on microbial community structure of *S. breviflora*, we systematically compared microbial community diversity across different ecological niches. Alpha diversity analysis revealed that divergent microbial community responses to grazing across bulk soil, rhizosphere, and root endophytic compartment (**Figure 2a-c**). Light grazing induced minor, non-significant fluctuations in Shannon diversity across all niches. Moderate grazing triggered niche-dependent divergent changes: microbial alpha diversity increased in bulk soil but decreased significantly in the rhizosphere and root endophytic compartment. Heavy grazing elevated Shannon diversity across all micro-niches, with a statistically significant increase exclusively in the rhizosphere.

**Figure 2.**
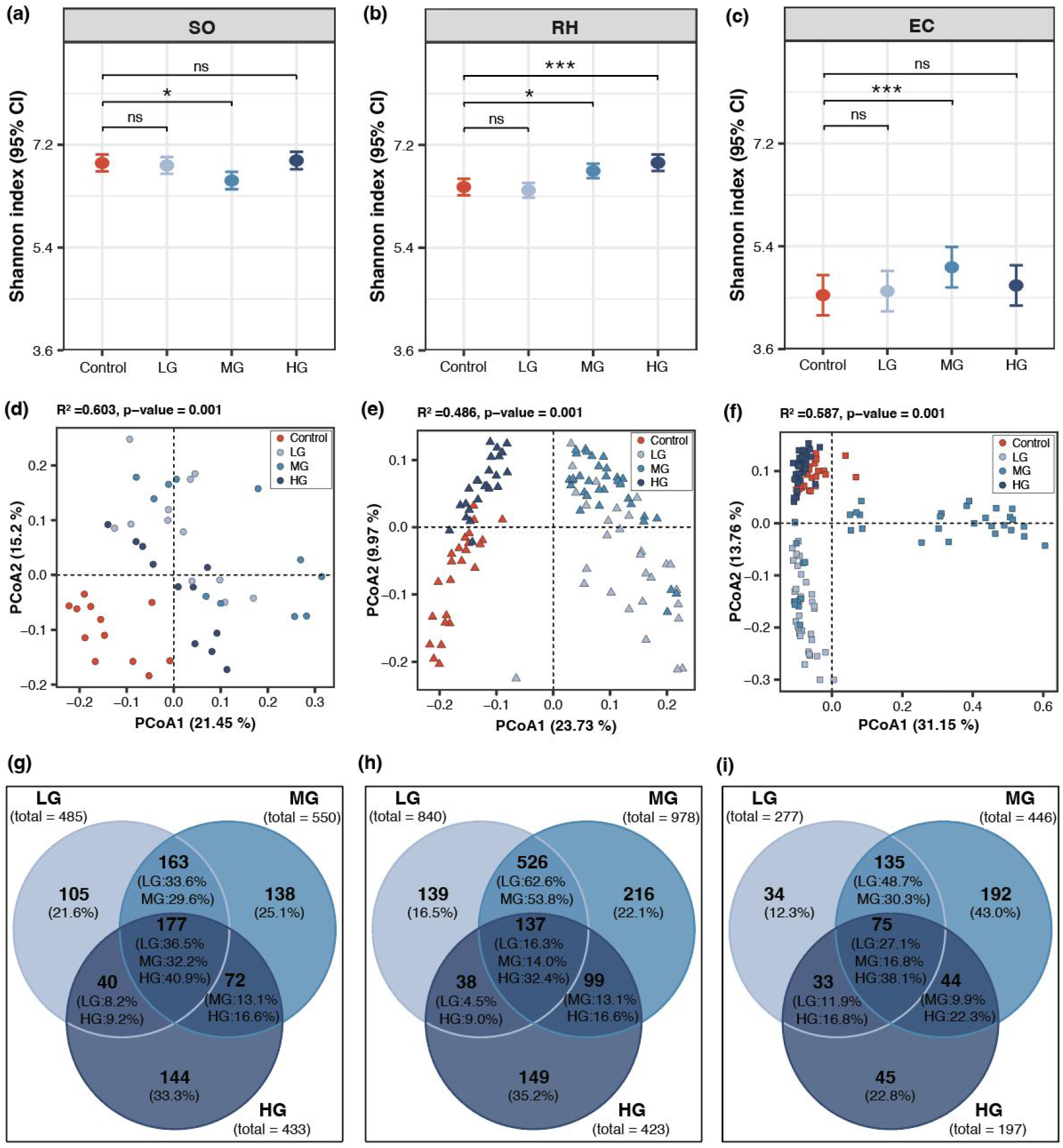
Grazing intensity affects microbial community structures. Grazing intensity affects the alpha-diversity of microbial communities in the bulk soil (a), rhizosphere soil (b), and root endosphere (c). Model-estimated means (bars) and 95% confidence intervals (error bars) of the Shannon index are shown for each grazing level. Principal Coordinates Analysis (PCoA) based on Bray-Curtis dissimilarity show differences in microbial community composition under different grazing intensities in the bulk soil (d), rhizosphere soil (e), and root endosphere (f). PERMANOVA (Adonis, transformed data by Bray-Curtis, permutation = 999) was used to determine the significant difference between the two groups. The Venn plots depict the shared number of the significantly altered ASVs in the bulk soil (g), rhizosphere soil (h), and root endosphere (i). Control, no grazing; LG, light grazing treatment; MG, moderate grazing treatment; HG, heavy grazing treatment. SO: Bulk soil, RH: Rhizosphere soil, EC: root endosphere. ns = not significant, * *= P*-value < 0.05, ** = *P*-value < 0.01, *** = *P*-value < 0.001.

Overall, moderate grazing exerted the most pervasive and substantial effects on microbial alpha diversity, while heavy grazing effects were restricted primarily to the rhizosphere.

Then, beta diversity analysis was performed with principal coordinate analysis (PCoA), which revealed distinct clustering of microbial communities under grazing, with the degree of separation varying by grazing intensity and compartments (**Figure 2d-f**). Moderate grazing samples exhibited clear segregation from control samples across all three niches with minimal overlap, representing the most pronounced community shift. Light grazing induced moderate community divergence with partial overlap with controls. In contrast, heavy grazing microbial communities displayed high similarity to the ungrazed control, with extensive axis overlap, particularly in the root endophytic compartment.

To further assess the impact of grazing intensity on microbial communities, we identified significantly differential ASVs (*P*-value < 0.05). Differential ASV analysis further verified the unimodal response of microbial communities to grazing disturbance (**Figure 2g–i**). Moderate grazing harbored the highest number of significantly altered ASVs (446–978), followed by light grazing (277–840), while heavy grazing induced the fewest microbial community changes (197–433). The number of differential ASVs under heavy grazing was less than half of that under moderate grazing in rhizosphere and endophytic compartment. The highest ASV overlap was observed between light and moderate grazing treatments (47.1%–78.9%), confirming that intermediate grazing disturbance is the primary driver of root-associated microbial community differentiation.

### Moderate grazing optimizes microbial co-occurrence network complexity and stability

To explore how grazing intensity reshapes the assembly and interaction patterns of soil microbial communities, microbial co-occurrence networks were constructed to characterize grazing-induced changes in microbial interspecific interactions based on significant correlations (|r| > 0.6, p < 0.001) among ASVs (**Figure 3**). The control network consisted of 489 nodes and 2,480 edges, with an average degree of 10.14 and a modularity of 0.477. Positive edges accounted for 66.73% of all connections, while negative edges accounted for 33.27%. Compared with the control, light grazing (LG) increased the number of nodes to 669 and the average degree to 12.38, but decreased modularity slightly to 0.459 and the proportion of positive edges to 55.24%. In contrast, heavy grazing (HG) reduced node number to 467 and average degree to 11.19, with a modularity of 0.450 and 64.32% positive edges, suggesting a simplification of network complexity under high grazing pressure.

**Figure 3.**
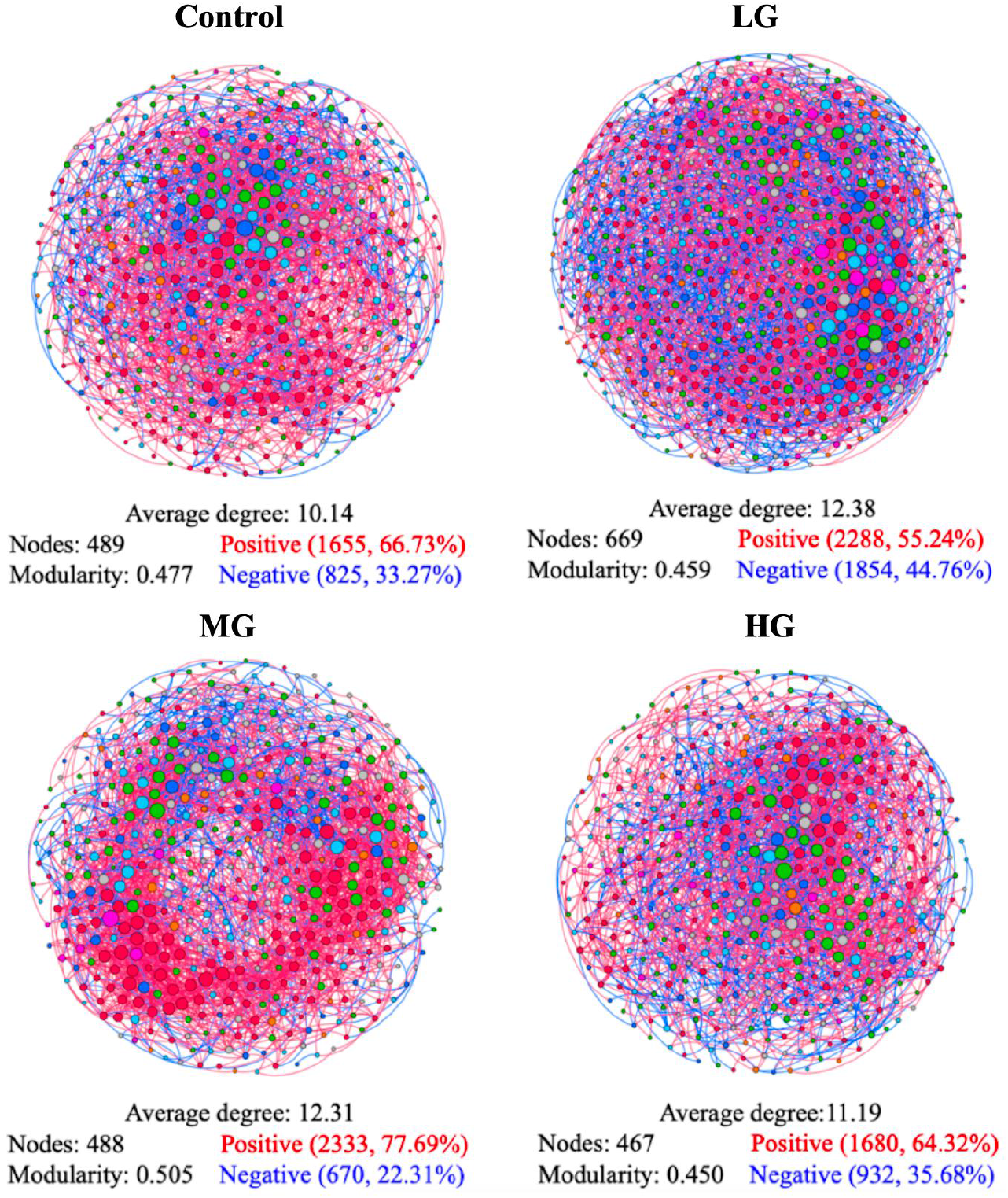
Microbial co-occurrence networks under different grazing intensities. Networks were constructed based on significant correlations (|r| > 0.6, p < 0.001) of ASV abundances. Each node represents an ASV, and each edge represents a significant co-occurrence relationship. Positive correlations are shown in red, and negative correlations in blue. Node size is proportional to degree (number of connections). Four treatments are presented: Control (no grazing), light grazing (LG), moderate grazing (MG), and heavy grazing (HG).

Notably, moderate grazing (MG) exhibited a distinct network structure compared to both light and heavy grazing. Although the node number in MG (488) was nearly identical to that of the control (489), the average degree increased markedly to 12.31, and modularity reached the highest value among all treatments (0.505). Furthermore, MG showed the highest proportion of positive edges (77.69%) and the lowest proportion of negative edges (22.31%). These results indicate that moderate grazing enhances cooperative microbial interactions, optimizes network modularity, and improves the stability and adaptive capacity of root-associated microbial communities.

### Grazing selectively enriches Bacillota and depletes *Pseudomonas* across micro-niches

To further investigate the specific microbial taxa affected by different grazing intensities, we performed a taxonomic composition analysis on the ASVs. Taxonomic composition analysis revealed consistent dominant bacterial phyla (Actinobacteriota, Proteobacteria, Acidobacteriota) in bulk and rhizosphere soils across all grazing treatments (**Figure 4a–c**). In the endophytic compartment, Actinobacteriota and Proteobacteria remained dominant, while light and moderate grazing specifically enriched Bacillota. Quantitative analysis confirmed that grazing significantly increased Bacillota relative abundance in a niche-dependent manner (**Figure 4d–f**). From bulk soil to rhizosphere and endophytic compartment, Bacillota abundance increased progressively, with significant enrichment in rhizosphere and endophytic compartments under light and moderate grazing; no significant variation was observed in bulk soil.

**Figure 4.**
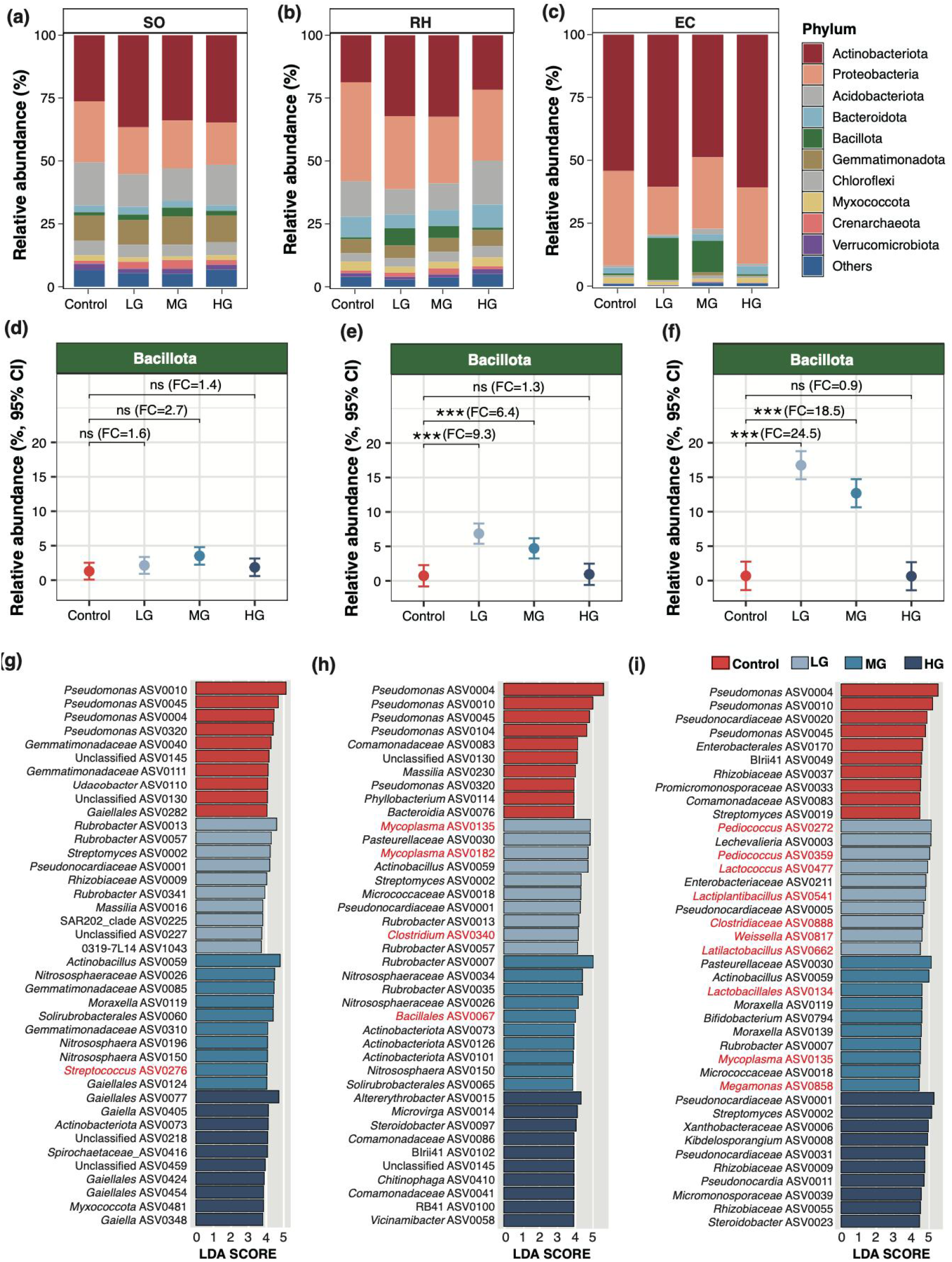
Changes in microbial taxa across grazing densities. Plots (a, b, c) display the relative abundance of the 10 most dominant bacterial phyla under different grazing densities in SO (a), RH (b), and EC (c). Control, no grazing; LG, light grazing intensity; MG, moderate grazing intensity; HG, high grazing intensity. Plots (d, e, f) highlight the relative abundance of the phylum Bacillota in SO (d), RH (e), and EC (f). Statistical significance in Bacillota abundance is indicated as: ns = not significant, ** = *p* < 0.01, *** = *p* < 0.001. FC, fold change. LEfSe plots showed the differential enrichment of ASVs in in SO (g), RH (h), and EC (i). The ASVs marked with red color belonged to the phylum Bacillota. Enrichment patterns were identified using linear discriminant analysis effect size with LDA score > 2.0.

To identify statistically significant and biologically relevant biomarkers (microbial taxa), Linear Discriminant Analysis Effect Size (LEfSe) analysis was performed. .LEfSe analysis revealed that Bacillota biomarkers increased progressively from bulk soil to endophytic compartment under light and moderate grazing treatments (**Figure 4g-i**). In bulk soil, only *Streptococcus* ASV0276 was enriched (moderate grazing). In rhizosphere, light grazing enriched three Bacillota biomarkers (*Mycoplasma* ASV0035, ASV0182, and *Clostridium* ASV0340), while moderate grazing enriched only *Bacillales* ASV0067. The most pronounced enrichment occurred in endophytic compartment: under light grazing, Bacillota accounted for 7 of the top 10 biomarkers (e.g., *Pediococcus* ASV0272, ASV0359, *Lactococcus* ASV0477); under moderate grazing, enriched Bacillota included *Lactobacillales* ASV0134, *Mycoplasma* ASV0135, and *Megamonas* ASV0858. Notably, *Pseudomonas* ASVs were consistently enriched across all niches in the control group but were significantly reduced by grazing treatments. Overall, grazing treatments shifted the biomarker profile from *Pseudomonas*-dominated to Bacillota genus-dominated, with the effect being most pronounced in the endophytic compartment under light grazing.

### Niche-specific enrichment of core Bacillota taxa under grazing disturbance

To further determine the taxonomic composition of grazing-responsive Bacillota ASVs, we extracted all ASVs classified within this phylum and identified 212 Bacillota ASVs primarily affiliated with *Lachnospiraceae, Lactobacillaceae, Clostridiaceae*, and *Ruminococcaceae* (**Figure 5**).

**Figure 5.**
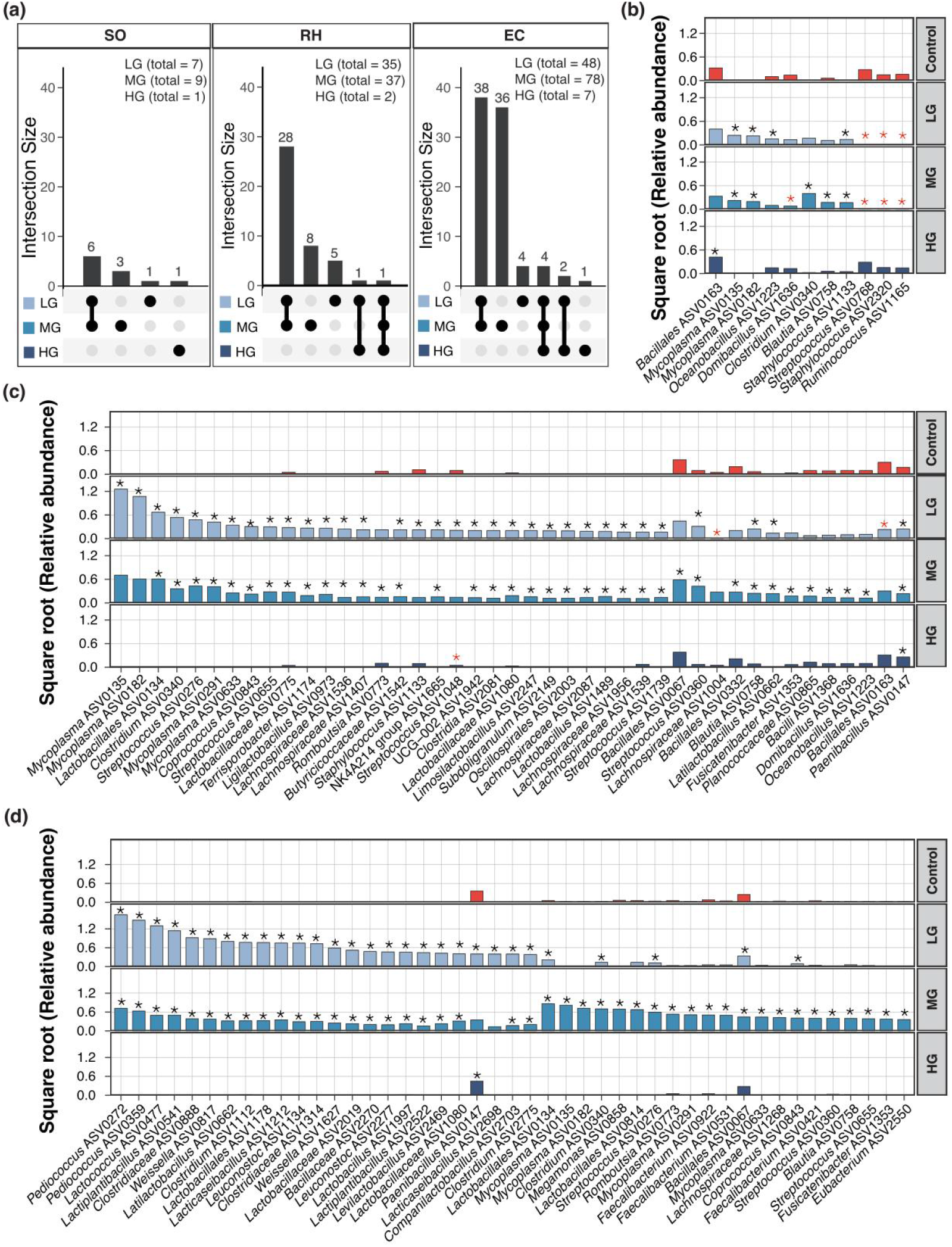
The impact of different grazing intensities on the microbes of the phylum Bacillota. (a) The Upset plots depict the number of overlapped microbial species that were significantly altered (increased or decreased) under various grazing density treatments when compared to the control. Control, no grazing; LG, light grazing intensity; MG, moderate grazing intensity; HG, high grazing intensity. Panels show the results derived from bulk soil (SO), rhizosphere soil (RH), and root endosphere (EC). Plots (b-d) detail the taxonomic composition and the relative abundance of the significantly changed microorganism in bulk soil (b), rhizosphere soil (c), and root endosphere (d). The bar height represents the square-root transformed relative abundance. Black asterisks denote a significant increase, while red asterisks indicate a significant decrease. Owing to the high number of significant changes in EC, the figure (d) displays a subset of the top 45 ASVs ranking highest in abundance.

Then, all Bacillota ASVs from each treatment group were used for further comparative analysis, and only a limited number of ASVs exhibited a significant decrease in relative abundance (compared to control). The number of significantly enriched Bacillota ASVs increased sequentially from bulk soil to rhizosphere and endophytic compartment, with moderate grazing inducing the strongest enrichment. Taxonomic profiling revealed niche-specific enrichment patterns: bulk soil was dominated by *Mycoplasma* (e.g., ASV0135, ASV0182) under light and moderate grazing; rhizosphere soil was enriched with *Mycoplasma, Streptococcus* and *Lachnospiraceae* under light grazing; the endophytic compartment exhibited unique *Pediococcus* enrichment under light grazing and dominant *Mycoplasma* enrichment under moderate grazing. Quantitatively, enriched Bacillota ASVs contributed 77.4% and 50.6% of total Bacillota abundance in the rhizosphere, and 98.8% and 88.6% in the endophytic compartment under light and moderate grazing, respectively. These results demonstrate that grazing intensity and soil-root micro-niche jointly determine the assembly and enrichment of stress-tolerant Bacillota communities.

## Discussion

Our 17-year field experiment demonstrates that long-term grazing drives dose-dependent and niche-specific restructuring of *S. breviflora* root-associated microbiomes, providing robust empirical evidence for microbial community adaptation to chronic anthropogenic activities in desert steppe ecosystems. The intermediate disturbance hypothesis (IDH) predicts that species diversity is highest at intermediate disturbance intensities, as no or low disturbance allows competitive dominance of species reducing diversity, whereas high disturbance imposes strong environmental filtering which also negatively affects diversity (Connell, 1978). In line with this hypothesis, we found that both light and moderate grazing maximized microbial community differentiation and biomarker abundance, while heavy grazing and a lack of grazing reduced microbial community variability and selective enrichment capacity. This unimodal response is most pronounced in rhizosphere and endophytic compartments, indicating that root exudation and host-mediated microbial recruitment amplify microbial sensitivity to grazing disturbance, whereas heavy grazing homogenizes belowground microbial assemblages.

Our taxonomic profiling identifies Bacillota and *Pseudomonas* as complementary microbial bioindicators for grazing disturbance. The significant enrichment of Bacillota (including *Paenibacillus, Clostridium, Lactobacillus*, and *Streptococcus*) under intermediate grazing is ecologically adaptive. These spore-forming taxa possess strong stress resistance, antimicrobial activity, and auxin synthesis capacity, enabling them to tolerate periodic plant defoliation, soil disturbance, and nutrient pulse fluctuations induced by livestock activity. The progressive enrichment of Bacillota from bulk soil to the endophytic compartment confirms active host-mediated recruitment, which likely alleviates grazing-induced oxidative stress and sustains plant nutrient uptake under disturbed conditions (Li et al., 2025). In contrast, *Pseudomonas*—canonical copiotrophic plant-growth-promoting rhizobacteria dependent on stable soil structure and high root exudation—dominated undisturbed grassland but was depleted under grazing. This distinct taxonomic trade-off reveals a functional shift of root microbiomes from “growth-promoting” to “stress-resilient” states under grazing disturbance.

From an ecosystem management perspective, our results demonstrate that both heavy overgrazing and long-term grazing exclusion induce functional simplification of root-associated microbiomes. Heavy grazing reduces microbial community differentiation and stress-tolerant taxon enrichment, while grazing exclusion locks the microbiome in a growth-promoting but low-resilience state dominated by *Pseudomonas*. We therefore propose that maintaining intermediate grazing disturbance via rotational or seasonal grazing regimes is critical for preserving microbial community diversity and ecosystem resilience in desert steppes. Future multi-omics integration (metagenomics and metabolomics) and microbial isolate inoculation experiments will further resolve the molecular mechanisms underlying Bacillota recruitment and their functional roles in plant drought and herbivory tolerance, supporting microbiome-targeted restoration of degraded grasslands.

## Conclusion

Long-term grazing functions as a selective ecological filter that profoundly restructures the diversity, composition, and interaction networks of desert steppe plant root-associated microbiomes in a niche- and intensity-dependent manner. Light and moderate grazing selectively enriches stress-tolerant Bacillota taxa while maintaining microbial community heterogeneity and network stability. By contrast, heavy grazing and grazing exclusion trigger microbial functional simplification and reduced ecosystem adaptive resilience. The contrasting responses of Bacillota and *Pseudomonas* to grazing disturbance establish them as reliable complementary bioindicators for desert steppe health monitoring. Integrating moderate disturbance grazing regimes with microbiome-based ecological interventions provides a promising strategy for sustaining the productivity and long-term resilience of arid and semi-arid grassland ecosystems.

## Notes

### Competing Interest Statement

The authors have declared no competing interest.

### Funding

- RZ2300003692 High-end Talent Acquisition Fund
- Innovation Team for Research on Interactions Between Grassland Plants and Microorganisms
- The “Golden Grain Talent” cadre of high-level professionals

